# Nrf1D is the first candidate secretory transcription factor in the blood plasma, with its precursor existing as a unique redox-sensitive transmembrane CNC-bZIP protein in somatic tissues

**DOI:** 10.1101/342428

**Authors:** Jianxin Yuan, Hongxia Wang, Shaojun Li, Yuancai Xiang, Shaofan Hu, Meng Wang, Lu Qiu, Yiguo Zhang

## Abstract

Amongst multiple distinct isoforms, Nrf1D is synthesized from translation of an alternatively-spliced transcript of Nrf1 mRNA, with a naturally-occurring deletion of its stop codon-flanking 1466 nucleotides. This molecular event leads to the reading frameshift mutation, which results in a constitutive substitution of the intact Nrf1’s C-terminal 72 amino acids (aa, covering the second half of the leucine zipper motif to C-terminal Neh3L domain) by an additional extended 80-aa stretch to generate a unique variant Nrf1D. The C-terminal extra 80-aa region of Nrf1D was identified to fold into a redox-sensitive transmembrane domain that enables it to be tightly integrated within the endoplasmic reticulum (ER) membranes. Notably, the salient feature of Nrf1D confers it to be distinguishable from prototypic Nrf1, such that Nrf1D is endowed with only a less ability than wild-type Nrf1 at mediating target gene expression. Further evidence has been presented revealing that both mRNA and protein levels of Nrf1D were detected to varying extents in somatic tissues. Surprisingly, we also found the existence of Nrf1D-derived isoforms in the blood plasma, implying that it is a candidate secretory transcription factor, although its precursor acts as an integral transmembrane-bound CNC-bZIP protein that entails dynamic topologies, before being unleashed from the ER to enter the blood plasma.

## INTRODUCTION

In the mid-1990s, nuclear factor-erythroid 2 (EF-E2) p45 subunit-related factor 1 (Nrf1, also designated as Nfe2L1) was identified as a key membrane of the cap’n’collar (CNC) basic-region leucine zipper (bZIP) family, which comprises Nrf2, Nrf3, Bach1, Bach2, Skn-1 and Cnc, in addition to p45 and Nrf1 [1-4]. Amongst this CNC-bZIP family, Nrf1 and Nrf2 are two principal master regulators of cognate target genes in mammals and also widely expressed in various distinct tissues. Later, ever-increasing experimental evidence has demonstrated that Nrf1 exerts its unique functions, which are distinctive from Nrf2 [5]. This notion has been supported by the fact that Nrf1, but not Nrf2, is indispensable for maintaining cellular homoeostasis and organ integrity during normal development and growth, albeit both factors are required for diverse adaptive responses to a variety of intracellular and environmental stresses, which are also involved in most pathophysiological processes [6, 7]. It is important to note that the essential function of Nrf1 is finely tuned by a steady-state balance of between its production and concomitant processing through post-transcriptional and post-translational pathways before being turned over. The processing of Nrf1 leads ultimately to generating various lengths of mRNA transcripts and protein isoforms (with different and even opposing abilities)[8, 9]. Overall, distinct Nrf1 isoforms are postulated together to confer on the host robust cytoprotection against cellular stress through coordinated regulation of distinct subsets of genes. Their transcriptional expression is predominantly driven by binding of Nrf1 to antioxidant response elements (AREs) or other homologous consensus sequences (i.e. AP-1 binding site) in those gene promoter regions.

In early studies using the consensus NF-E2/AP1-binding sites as a probe to clone the cDNA sequence of Nrf1, it was identified to consist of 742 aa in human [1] or 741 aa in mouse [10]. Similar cloning strategies were also employed to identify LCR-F1 [4] and TCF11 [2] that comprise 447 and 772 aa, respectively. Except length variations, both the nucleotide and amino acid sequences of LCR-F1 and TCF11 are identical with equivalents of Nrf1, and thus they are thought as different length isoforms [5]. In fact, the prototypic Nrf1 (i.e. its full-length protein Nrf1x) is generated by translation from alternative splicing of mRNA to remove exon 4 [that encodes ^242^VPSGEDQTALSLEECLRLLEATCPFGENAE^271^, called Neh4L region] from human TCF11 [2]. As such Neh4L is lost in Nrf1, it was shown to exhibit similar transactivation activity to that of TCF11 [11], but this long TCF11 is not found in the mouse [10]. In addition, the post-synthetic processing of Nrf1/TCF11 may also yield multiple distinct polypeptides of between 140-kDa and 25-kDa, which together determine its overall activity to differentially regulate target genes [8, 9, 12]. Further comparison of amino acid sequences demonstrates that LCR-F1 is a shorter form of Nrf1 (i.e. Nrf1β) [13], which is translated by its in-frame perfect Kozak initiation signal (5’-puCCATGG-3’) that exists around the methionine codons at between positions 289-297 in the mouse [1, 2, 10]. Relative to Nrf1, Nrf1β/LCR-F1 lacks the N-terminal acidic domain 1 (AD1) [11, 14] and hence exhibits only a weak transactivation activity [4, 8, 15, 16]. As such, Nrf1β/LCR-F1 activity may also be differentially induced in responses to distinct stressors [15-17]. In addition, Nrf1β/LCR-F1 is unstable because it may be rapidly processed to yield two small poplypeptides of 36-kDa Nrf1y and 25-kDa Nrf18 [8, 9]; both may also be generated by additional in-frame translation. Importantly, these two small dominant-negative Nrf1y and Nrf18, when over-expressed, have a capability to competitively interfere with the functional assembly of the active CNC-bZIP transcription factors, so as to down-regulate expression of NF-E2/AP1-like ARE-driven genes [8, 15].

Distinct other isoforms of Nrf1 have been determined to arise from multiple variants of mRNA transcripts, most of which are deposited in both GenBank and Ensembl (i.e. ENSMUSG00000038615 and ENSG00000082641, representing mouse and human *Nef2l1*, respectively). For example, those variants in the 3’- and 5′ -untranslated regions were found to yield four different types of mRNA transcripts, that are consequently translated into distinct lengths of Nrf1 isoforms with different trans-acting potentials [2]. Additional isoforms were also identified to be generated from alternative splicing of mRNA transcripts [15, 16]. Amongst these variants, two are the most peculiar, that were originally designated as Nrf1 clones Δ767 and D, with a deletion of its original translational start or stop codons, respectively. Removal of the first translation start codon results in production of an N-terminally-truncated isoform designated as Nrf1^ΔN^ [12, 13], within which the first N-terminal 181-aa region of Nrf1/TCF11 is replaced by an extra dodecylpeptide MGWESRLTAASA. By contrast, the variant clone D of Nrf1 does not only lack the intact original translation stop codon, but also acquires a substitutive change in the second half of its leucine zipper domain along with an extended C-terminal region, which is hence renamed Nrf1D (Fig. 1A). However, both the biochemical characteristics and biological behaviours of Nrf1D have remained elusive to date. To address these, we herein examine whether: i) it has an unique C-terminal transmembrane (TMc) region, which is distinctive from the counterpart of intact Nrf1, involved in topovectorial process; ii) the TMc enables this variant to become more sensitive to oxidants and reducing agents, when compared with wild-type protein; and iii) it is allowed for its putative processing to yield a putative specific isoform that is likely secreted in the blood plasma, albeit most of this variant exists widely in somatic tissues.

**Figure 1.**
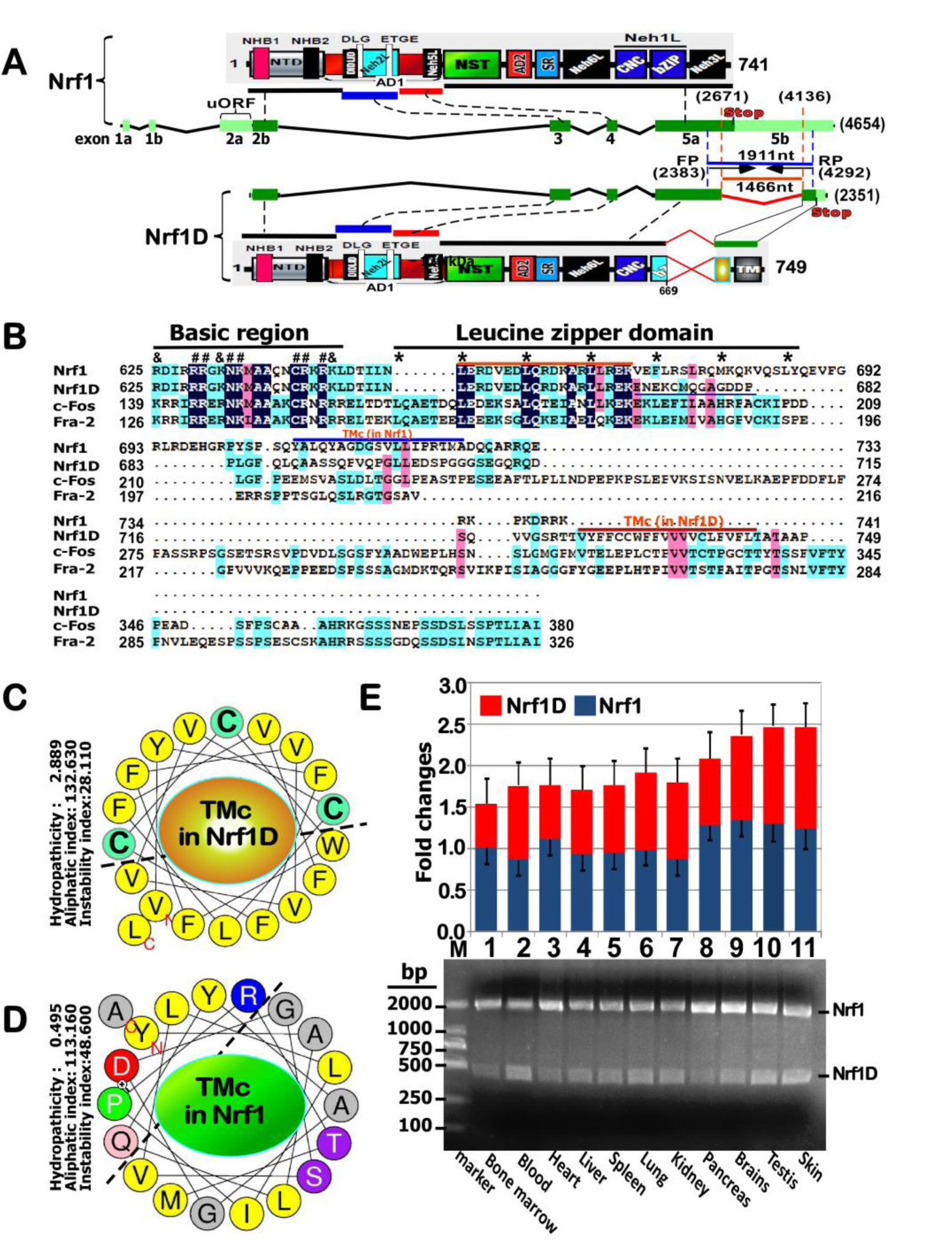
Structural differences in between Nrf1D and Nrf1 with distinct transcripts expressed in different tissues. (***A***) Schematic representation of structural domains of mouse Nrf1 and its splicing variant Nrf1D, which are translationally expressed by distinct exons of both transcripts, respectively. (***B***) Amino acid alignment of bZIP and C-terminal adjacent domains in Nrf1, Nrf1D, c-Fos and Fra-2. The basic region, leucine zipper domain, and TMc stretch are indicated. (***C, D***) Two different a-helical wheels folded by those amino acids covering the C-terminal transmembrane (TMc) domains in Nrf1D and Nrf1. Within TMc of Nrf1D, the thiol-based Cysteine residues are placed in green backgrounds, whilst hydrophobic and other aliphatic amino acid residues are in yellow backgrounds (*C*). (***E***) Differential expression of Nrf1 and Nrf1D transcripts in 11 different tissues from the mouse was determined by RT-PCR with a pair of specific forward and reverse primers described in the section of “Materials and Methods”. The data are here shown as a fold change (mean ± S.D), each of which represents at least 3 independent experiments undertaken on separate occasions that were each performed in triplicate. In addition, the fidelity of both Nrf1 and Nrf1D transcripts wad further confirmed by sequencing, as their nucleotide sequence alignment was shown in Fig. S1.

## RESULTS

### Nrf1D has a unique C-terminal 80-aa region with a potential redox-sensitive TMc fold

By bioinformatic analysis of both Nrf1D and Nrf1 (Fig. 1, A and B), it was revealed that the C-terminal 72-aa residues of wild-type Nrf1, which are extremely positively charged, are exchanged for additional 80-aa stretch, which is, however, enriched with negatively charged residues in the variant Nrf1. Further sequence alignment showed that the extra 80-aa region of Nrf1D, but not the equivalent sequence of Nrf1, appears to have a certain conservativity with c-Fos and Fos-related antigen-2 (Fra-2) (Fig. 1B). Nonetheless, Nrf1, rather than c-Fos or Fra-1, was also predicted to form a redox-sensitive (i.e. cystines at positions 729, 730 and 738) hydrophobic TMc a-helix folded on its C-terminal end (Fig. 1C). The wheeled TMc region of Nrf1D has acquired an enhanced hydropathicity of 2.889, which is 5.8 times higher than the counterpart value (0.495) of Nrf1, despite of no marked difference in both aliphaticity (*cf. panels C with D*). Thus, it is postulated that Nrf1D is also tightly integrated by its C-terminal TMc region spanning across within the endoplasmic reticulum (ER) membrane, whereas the equivalent of Nrf1 could be loosely tethered to membranes with dynamic topologies entailed. In addition, the commonly-sharing basic-region and the first half of the leucine zipper domain of Nrf1 and Nrf1 are highly conserved with those of c-Fos and Fra-2. Collectively, it is assumed that Nrf1D is better than Nrf1, acting as a substitute for c-Fos in the formation of a heterodimmer with the latter partner c-Jun.

### Nrf1D exists widely in bloods, bone marrows and somatic tissues examined

To investigate differences in mRNA expression levels of Nrf1D and Nrf1 in distinct tissues, semi-quantitative real-time PCR was monitored by an pair of optimal primers, that were complementary to their two commonly-sharing upstream and downstream nucleotide sequences indicated (Fig. 1A). These results clearly demonstrated the existence of the variant *Nrf1D* in all tissues examined (Fig. 1E). The expression profiles of *Nrf1D* mRNA were roughly similar to, and even higher than, the corresponding levels of wild-type *Nrf1*, particularly in mouse blood, lung, brains, testis and skin. All the short cDNA fragments of Nrf1D obtained from PCR products were validated by sequencing and also confirmed to be true by a multiple nucleotide sequence alignment (Fig. S1).

To determine distinction in their protein expression profiles of between Nrf1D and Nrf1, its specifically-recognized antibody was made herein and further verified by immunoblotting of total lysates of COS-1 cells that had been transfected with an expression construct for Nrf1D, Nrf1 or empty pcDNA3 vector. The results showed that expression of Nrf1 was identified by its unique immunoreactivity with the *per se* C-terminal peptide-specific antibody (Fig. 2B), even though it was also recognized by additional two antibodies against aa 192-741 of Nrf1 (Fig. 2A) and its C-terminal V5 tag (Fig. 2C), respectively. The mobility of Nrf1 polypeptides to electrophoretic locations of between 140-kDa and 25-kDa revealed that it appeared to have undergone the post-synthetic processing to give rise to its protein ladder, probably through a similar mechanism accounting for Nrf1, as consistent with the notion reported previously [9, 18].

**Figure 2.**
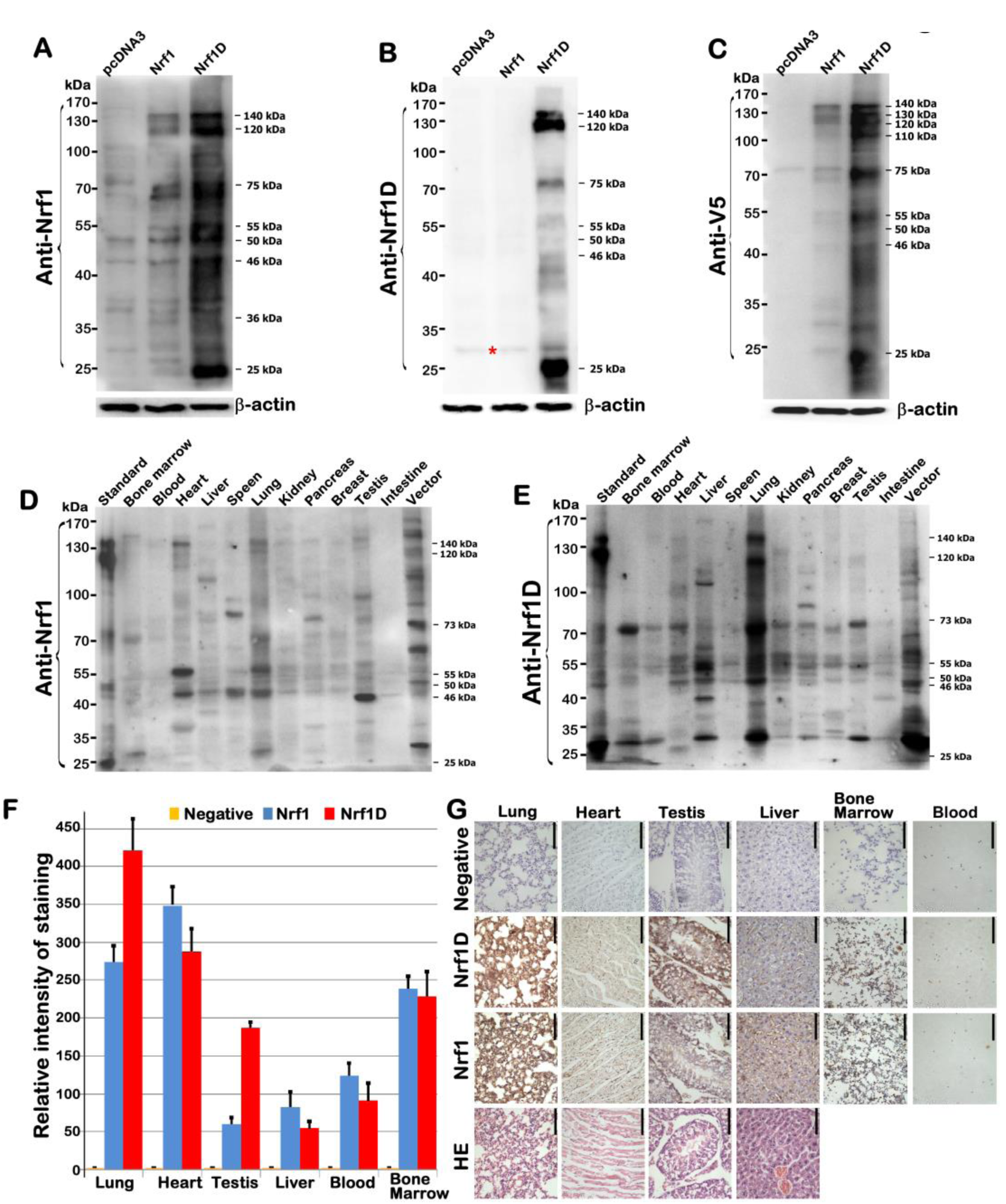
Differential expression of Nrf1D and Nrf1 proteins in mouse tissues examined. (***A*** *to**C***) Both Nrf1D and Nrf1 were identified by cross-immunoreactivity with three distinct antibodies against aa 192-741 in Nrf1, Nrf1D’s C-terminus-specific peptide, and their C-terminally-tagged V5 ectope, respectively. (***D,E***) Differential expression levels of Nrf1- and Nrf1D-derived proteins in different tissues from mice were examined by Western blotting with Nrf1- or Nrf1D-specific antibodies. (***F,G***) Anti-Nrf1D, and -Nrf1 antibodies were used in immunohistochemistry of six different tissues, including lung, heart, testis, liver, bone marrow and blood (bar = 100 µm). Subsequently, the intensity of immunohistochemical staining in examined tissues was calculated and is shown graphically (*G*).

Further immunoblotting of different tissues unveiled that differential expression profiles of endogenous Nrf1D (Fig. 2E) and Nrf1 (Fig. 2D) were exhibited with distinct lengths of isoforms, even in each of the same organs. The variations in the protein expression of Nrf1 were also detected in single individual animals with gender differences (Fig. S2). The full-length Nrf1D of ^~^140-kDa and its four major isoforms of ^~^115-, 70-, 45- and 35-kDa were determined primarily in the lung (Figs. 2E and S2). As such, some of Nrf1D and/or its derivates of between 115-kDa and 45-kDa were also, to greater or less extents, examined in other tissues, such as the blood, bone marrow, heart, liver, kidney, pancreas, brains and testis. By contrast, the wild-type Nrf1 of 140-kDa and its isoforms of between 90-kDa and 45-kDa were dominantly expressed in bone marrow, lung, heart and testis, apart from less abundances of them detected in other tissues (Fig. 2D). Further immunohistochemistry displayed distinctive staining with the Nrf1D-specific antibody, when compared with that obtained from anti-Nrf1 staining (Figs. 2F,G, and S3 to S5). This observation indicates differential expression of both CNC-bZIP proteins in the lung, heart, testis and liver, as well as in the bone marrow-derived and blood cells.

To clarify the presence of Nrf1D in the bone marrows, the fluorescence *in situ* hydridization (FISH) was employed with an Nrf1D-specific probe (5’-CGATTGCTTCGAGAAAAGGAAAATGAGAAGTGC-3’, with an high-fidelity to hybridize two small segments flanking its original mRNA alternatively-spliced and ligated sites). The resulting *in situ* hydridization of Nrf1D or Nrf1 mRNA transcripts by their cDNA probes was illustrated as red and green fluorescent signals, respectively (Fig. 3,B and C). The single signals of Nrf1D, Nrf1 and/or their superimposed signals, were displayed exclusively in the juxtanuclear locations of the bone marrow-derived cells (in particular megakaryocytes and polykaryocytes), albeit this is required for further studies in details.

**Figure 3.**
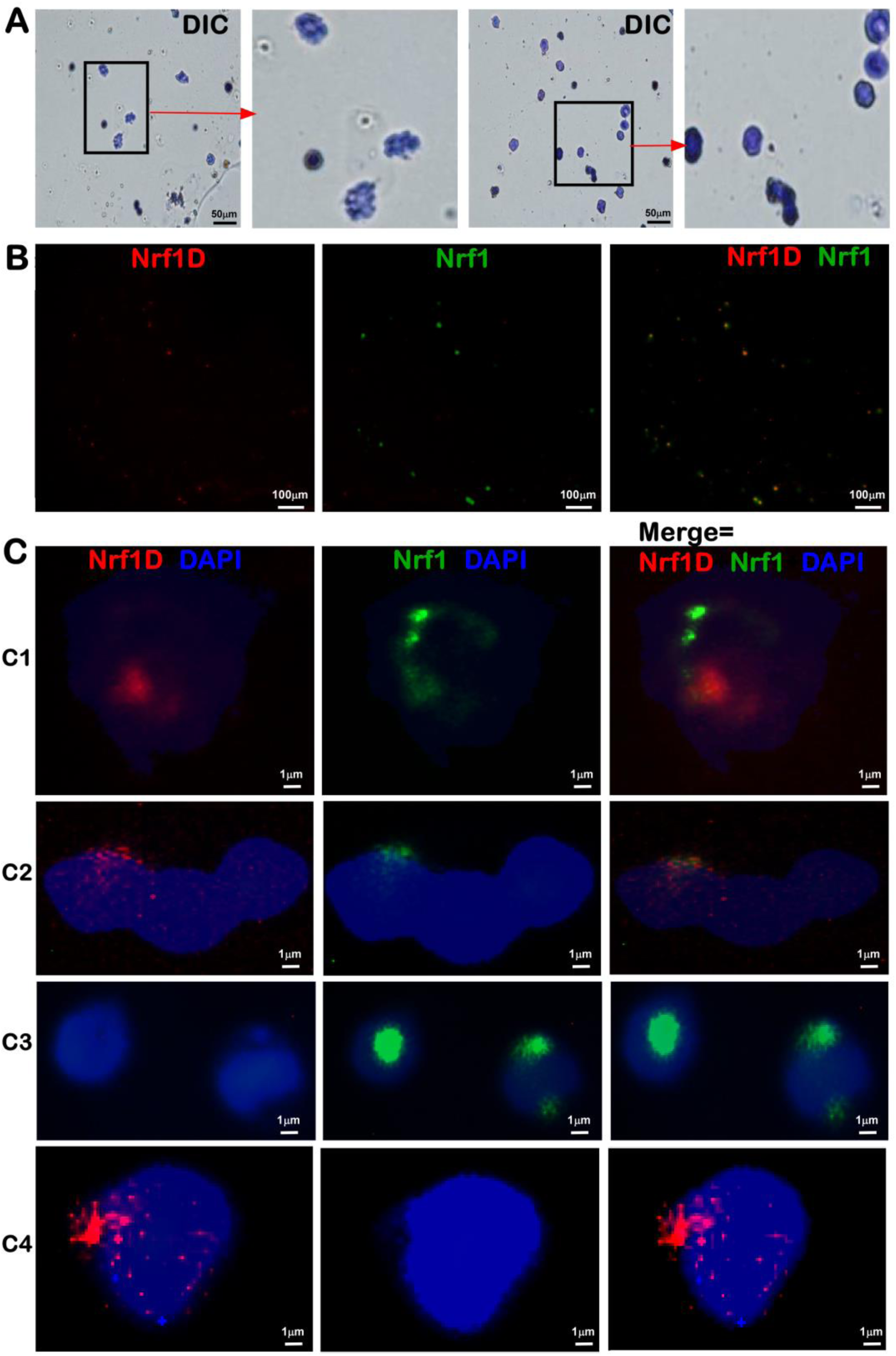
Fluorescence in situ hybridization (FISH) of Nrf1D and Nrf1 transcripts in bone marrow-derived cells. (***A***) Differential interference contrast (DIC) images (bar = 50 µm) of mouse bone marrow-derived cells were achieved in two different fields of vision by microscopy. (***B,C***) FISH of Nrf1D (red) and Nrf1 (green) transcripts expressed in bone marrow cells was conducted with their specific probes of Nrf1 (5’-CGCACGATGGCTGACCAGCAGGCTC-3’, which retained in Nrf1 but not Nrf1D, with its 5’-end labeled by 5-FAM, emitting a green fluorescence) and Nrf1D (5’-CGATTGCTTCGAGAAAAGGAAAATGAGAAGTGC-3’, which was labeled by Texas Red at its 5’-end so as to emit a red fluorescence). Nuclear DNA was stained by DAPI (blue). These images of indicated cells with different magnification (*B*, bar = 100 µm; *C*, bar = 1 µm) were acquired in different visual fields.

### Nrf1D is a candidate secretory transcription factor

Despite the existence of Nrf1D and Nrf1 in blood and hematopoietic cells, it is a curiosity that pushes us to examine whether both factors and/or their derivates come into emergence in the blood plasma. As excepted, the initial results unveiled an immunoreactivity of the plasma constituents (which also contained ^~^55-kDa IgG), rather than red blood cells (RBC), with antibodies against Nrf1 and Nrf1D (Fig. 4A). Further immunoblotting with the Nrf1D-specfic antibody revealed a major ^~^120-kDa protein along with an obvious polypeptide ladder between 90-kDa and 25-kDa in the upper plasma (Fig. 4B, *the second lane*), but a few of polypeptides between 70-kDa and 45-kDa were detected in the middle, rather than lower, fractions of blood obtained from male mice. By contrast, almost none of longer Nrf1D forms, except shorter polypeptides of between ^~^80-kDa and 45-kDa, were examined in those corresponding fractions of blood from female mice (Fig. 4B, *left three lanes*). Nevertheless, it is of significant importance to note that almost not any forms of Nrf1 were detected in the lower fractions (Fig. 4,A and B), which were enriched with erythrocytes (45% of total blood), at the bottom of the centrifuge tube, although the wild-type Nrf1 and its homologues were *ab initio* isolated in erythroid hematopoietic cells [1,2,4].

**Figure 4.**
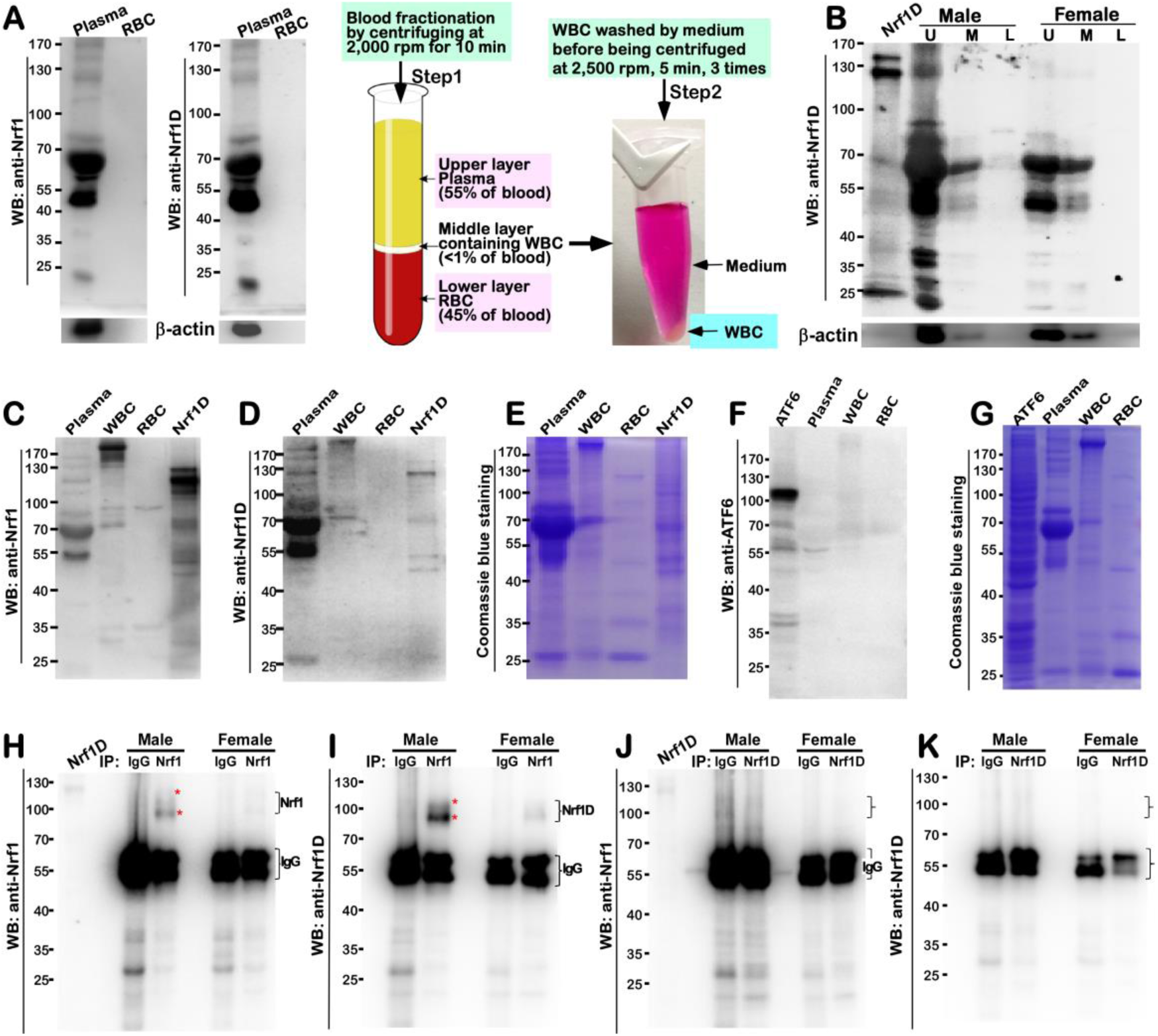
Detection of Nrf1D existing in mouse blood plasma. (***A***) Shows immunoblots of mouse blood plasma and RBC with anti-Nrf1 and Nrf1D-specific antibodies (*left upper two panels*). The procedure of blood fractionation by centrifuging at different speeds was illustrated (*right two panels*). (***B***) Distinct blood fractions from male and female mice were determined by Western blotting with Nrf1D-specific peptide antibody. The Nrf1D-expressed cell lysates served as a positive control. In addition, secreted β-actin was detected in blood plasma (*bottom*). Abbreviations: *U*, upper layer of plasma; *M*, middle layer containing WBCs; *L,* lower layer containing RBCs. (***C to G***) Further extracts from blood plasma, RBCs and WBCs were visualized by immunoblotting with Nrf1 antibody (***C***) or Nrf1D-specific peptide antibodies (***D***). However, similar blood fractions were not cross-immunoreactive with ATF6-specific antibodies (***F***). Total protein extracts from blood fractions were seen by Coomassie Blue staining (***E,F***). (***H to K***) Immunoprecipitation (IP) of blood plasma from male and female mice was employed with anti-Nrf1, anti-Nrf1D antibodies or IgG (served as an internal negative control). Subsequent immunocomplexes were visualized by Western blotting with indicated antibodies.

The IgG-depleted plasma was further subjected to immunoblotting with antibodies against Nrf1 (Fig. 4C) and Nrf1D (Fig. 4D). The results showed that Nrf1 and Nrf1D together with their derivates were migrated by electrophoresis to locations between 140-kDa and 45-kDa enriched in the plasma (*the left first lane*), when compared with the positive expression controls (i.e. *the lane labeled by Nrf1D*). However, almost none of longer Nrf1 proteins between 140-kDa and 100-kDa) were detected in the purified fractions of WBCs (i.e. leukocytes) and RBCs (i.e. erythrocytes), albeit with an exception that only a few of the minor polypeptides of between 90-kDa and 70-kDa were determined to have less immunoreactivity with antibodies against Nrf1 (Fig. 4C), but not Nrf1D-specfic peptide (Fig. 4D). In addition, a possible non-specific immunoreactivity appeared to be ruled out by the parallel experiments, which demonstrated that not an additional transmembrane-bound ATF6 and processed polypeptides existed in the blood plasma (Fig. 4F,G).

To further verify the existence of Nrf1 and/or Nrf1 in the blood plasma, their antibodies were utilized in distinct experimental settings of immunoprecipitation (IP). The resulting immunoprecipitates of between ^~^110-kDa and 95-kDa with Nrf1 antibody were not only recognized by this antibody *per se* (Fig. 4H) and also visualized by Western blotting (WB) with Nrf1D-specific antibody (Fig. 4I). However, it is regretful that none of these precipitates were obtained from Nrf1D-peptide antibody, because they were not detected by Western blotting with antibodies against Nrf1 (Fig. 4J) or Nrf1 antibody itself (Fig. 4K). Collectively, these results suggest a possibility that the antigen-specific peptide of Nrf1 is much likely to be buried in the interior of certain folding conformation such that it cannot be exposed, hence leading to no access to cognate antibodies during immunoprecipitation. This notion appears to be supported by the finding that Nrf1D-expressing cell lysates served as a positive control, with a strong cross-reactivity with Nrf1 antibody, but only a weak cross-reactivity was measured by Nrf1D-specific antibody (Fig. 4, C & D). Rather, additional possibility cannot be ruled out that the putative processing of Nrf1D might also result in proteolytic removal of its C-terminal peptides from this protein so that none of the immunoprecipitates were captured. Moreover, the immunoprecipitates with Nrf1 antibody were pulled down principally form male mice, but less or none of them were observed in female mice (Fig. 4,H and I), implying a possible gender difference.

### Dynamic topovectorial moving of Nrf1D in and out of the ER is also determined by its unique TMc region

Clearly, all those known secretory and transmembrane proteins as elucidated in the current literature [19, 20], are biosynthesized by ribosomes budded with the ER. Therefore, we examined whether the membrane-topological folding of Nrf1 is also determined by its unique TMc region, which is distinctive from the equivalent of Nrf1 that is integrated within and around membranes [9, 18, 21] (Fig. 5B1). To address this, the live-cell imaging of Nrf1D-eGFP (Fig. 5A) and Nrf1D^ΔDC^-eGFP (lacking the unique C-terminal 80-aa region of Nrf1D, Fig. S6A) in combination with *in vivo* membrane protease protection assays, was performed to clarify whether these two fusion proteins have capability of being released from the ER into extra-ER compartments. COS-1 cells expressing Nrf1D-eGFP or Nrf1D^ΔDC^-eGFP, together with the ER/DsRed marker, were pre-treated for 10 min with digitonin (to permeabilize cellular membranes) before being challenged with proteinase K (PK) for 3-30 min (still in the presence of digitonin) to digest cytoplasmic proteins. Hence, we surmised that if these two fusion proteins were transferred from the ER luminal side of the membrane into the cytoplasmic side and also dislocated into extra-ER compartments, it would become vulnerable to digestion by PK. As anticipated, the green fluorescent signals from Nrf1D-eGFP (Fig. 5A) and Nrf1D^ΔDC^-eGFP (Fig. S6A) were superimposed upon the red fluorescent images presented by ER/DsRed, implying that both fusion proteins are localized in the ER.

**Figure 5.**
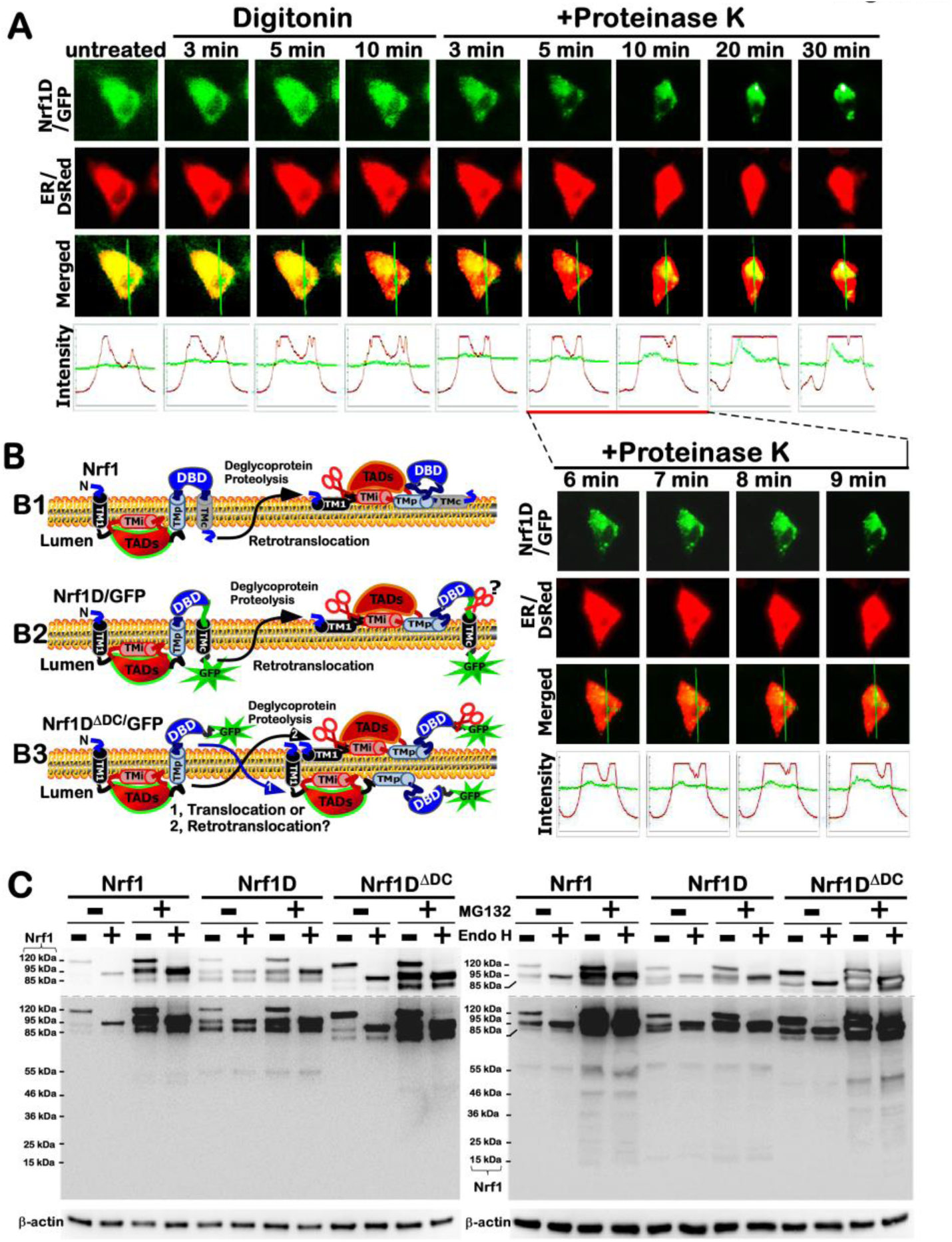
Dynamic repositioning of Nrf1D out of the ER membrane. (***A***) COS-1 cells that had been co-transfected with expression constructs for Nrf1D-eGFP and the ER/DsRed marker were subjected to live-cell imaging combined with *in vivo* membrane protease protection assay. The cells were permeabilized by digitonin (20 μg/ml) for 10 min, before being co-incubated with PK (50 μg/ml) for additional 30 min. During the course of the experiment, real-time images were acquired using the Leica DMI-6000 microscopy system. The merged images of Nrf1-eGFP with ER/DsRed are placed (*on the third row of panels*), whereas changes in the intensity of their signals are shown graphically (*bottom*). Overall, the images shown are a representative of at least 3 independent experiments undertaken on separate occasions. (***B***) Distinct membrane-topologies of Nrf1, Nrf1D-GFP, and Nrf1D^ΔDC^-GFP are depicted schematically. Transactivation domains (TADs), DNA-binding domain (DBD), and putative membrane-bound regions such as TM1, TMi, TMp and TMc are indicated. (***C***) Total lysates of COS-1 cells that had been transfected with expression constructs for Nrf1, Nrf1D or Nrf1D^ΔDC^ and treated for 4 h with 5 µmol/L MG132 were subjected to *in vitro* deglycosylation reactions with Endo H (+) or without this enzyme (−) for additional 4 h at 37°C. These reaction products were resolved by gradient LDS-NuPAGE gels containing 4-12% polyacrylamide and visualized by Western blotting with V5 antibody

When compared to untreated cell status, almost no changes in the intensity of the green signals from Nrf1D-eGFP were observed within 10 min after treatment of cells with digitonin and even in the presence of PK for 3 min (Fig. 5A). This demonstrates convincingly that the GFP ectope attached to the C-terminus of Nrf1 was initially translocated into the ER and buried in the lumen (Fig.5B2), so that it was protected by the membrane against PK digestion. As the time of additional PK treatment was further extended from 5 min to 30 min, the green signals from Nrf1D-eGFP became gradually fainter, but did not disappear until the experiment was stopped (Fig. 5A). By contrast, the ER/DsRed signals were not only reduced by PK digestion, but also obviously enhanced; this was attributable to that the cells were shrunken in sizes during this treatment. Collectively, these observations indicate that a large fraction of Nrf1D-eGFP is dynamically repositioned from the ER lumen into extra-ER subcellular compartments (i.e. cyto/nucleoplasm), but another small fraction of this fusion protein is tightly protected by ER membranes against PK digestion (Fig. 5B2).

By comparison with Nrf1D-eGFP, it is interesting to note that the green signals from Nrf1D^ΔDC^-eGFP were rapidly reduced following treatment of the cells with digitonin alone, and further diminished by additional PK digestion of this fusion protein for 3 to 10 min, albeit the time-lapsed signals were not eliminated by PK until the digestion time was extended to 30 min(Fig. S6A). This observation, together with our previous work [9, 18], indicates that a major fraction of Nrf1D^ΔDC^-eGFP (lacking the TMc region) was processed in the ER so as to be rapidly dislocated and released into extra-ER extracellular compartments after cellular membrane was permeabilized by digitonin, but another small fraction of this mutant fusion protein was retained in the ER lumen and thus protected by membranes (Fig. 5B3). Collectively, the subtle nuance in the membrane-topological folding of between Nrf1D-eGFP and Nrf1D^ΔDC^-eGFP suggests that dynamic moving of Nrf1 in and out of the ER is also determined by its unique TMc region, beyond its N-terminal TM1 region as described in wild-type Nrf1.

### Post-translational processing of Nrf1D to yield distinct isoforms in and out of the ER

*In vitro* endoglycosidase (Endo)-H reactions (Fig. 5C) revealed that, like wild-type Nrf1 as described [9, 18], Nrf1D was also N-glycosylated to become a ^~^120-kDa glycoprotein, which is then deglycosylated to yield another ^~^95-kDa deglycoprotein, prior to being proteolytically processed to generate an ^~^85-kDa N-terminally-truncated isoform. The selective proteolytic processing of Nrf1D, as well as its C-terminally-truncated mutant Nrf1D^ΔDC^, is postulated to have occurred by a potential mechanism similar to that accounting for Nrf1, even in the presence of proteasomal inhibitor MG132 (Figs. 5C & S7).

The pulse-chase experiments showed subtle differences in the conversion of distinct isoforms arising from amongst Nrf1, Nrf1D and Nrf1D^ΔDC^ and their stability before being turned over (Fig. S7). The half-lives of ^~^120-, 95- and 85-kDa isoforms yielded from Nrf1 were estimated to 0.33 h (= 20 min), 1.40 h (= 84 min) and 1.50 h (= 90 min), respectively (Fig. S7A), after Nrf1-expressing cells had been treated with 50 gg/ml cycloheximide (CHX, which inhibits biosynthesis of nascent proteins). Addition of MG132 caused a significant prolongation of the ^~^120-kDa Nrf1 glycoprotein’s half-life to 2.10 h (= 126 min), implying that this proteasomal inhibitor may have an effect on conversion of Nrf1 into the 95- and 85-kDa isoforms. As such, the half-lives of the 95- and 85-kDa Nrf1 isoforms were also concomitantly extended to over 8 h, suggesting both protein degradation by proteasome-mediated pathways.

By contrast with Nrf1 isoforms, the abundance of ^~^120-kDa Nrf1 glycoprotein was much less than those levels of its 95-kDa deglycoprotein and processed 85-kDa proteins (Fig. 7B, *left panel*), indicating that the ^~^120-kDa glycoprotein is rapidly converted into the 95-kDa deglycoprotein and processed 85-kDa isoforms. Such conversion of Nrf1D-derived isoforms and these protein stability were determined by distinctive extension of their half-lives to 0.88 h (= 53 min), 2.46 h (= 148 min) and 6.44 h (= 386 min) respectively, after CHX treatment (Fig. S7B), implying that Nrf1D-derived proteins are more stable than those equivalents of Nrf1. Intriguingly, further examinations revealed that abundances of Nrf1D-derived ^~^120-, 95- and 85-kDa proteins were only marginally promoted by additional treatment of cells with MG132 (Fig. 7B, *left panel*), with modest alterations of their half-lives estimated to be 0.63 h (= 38 min), 3.13 h (= 188 min) and 7.71 h (= 463 min), respectively, after co-treatment with CHX (Fig. S7B, *right graphs*).

The above finding demonstrates that a considerable fraction of distinct Nrf1D-derived isoforms is irresponsive to the proteasome-mediated proteolysis. This fact suggests that they may also be targeted to a proteasome-independent degradation pathway, but it cannot be ruled out that Nrf1D-derived isoforms may be partially protected by its unique TMc-associated membranes against cytosolic proteases. To explore this possibility, the TMc-truncated Nrf1D^ΔDC^ mutant was subjected to the pulse-chase experiments. This mutant protein was N-glycosylated into an ^~^110-kDa glycoprotein, and then deglycosylated to yield a ^~^85-kDa deglycoprotein, before being proteolytically processed to give rise to an N-terminally-truncated ^~^75-kDa protein (Figs. 5C and S7C). The conversion of Nrf1D^ΔDC^ -derived ^~^110-, 85- and 75-kDa proteins and their stability were also determined with distinct half-lives, estimated to be 0.69 h (= 41 min), 0.53 h (= 32 min), and 1.00 h (= 60 min) respectively, after CHX treatment (Fig. S7C, *right graphs*). Their half-lives were significantly prolonged by additional treatment with MG132 to 2.16 h or more than 8 h, respectively. Collectively, together with the data as described above (Fig. S6A), these findings suggest that Nrf1 is tightly protected by its unique TMc-associated membranes, when compared with the TMc-truncated mutant Nrf1D^ΔDC^. It is inferable that the ^~^110-kD glycoprotein of Nrf1D^ΔDC^ is rapidly repositioned into extra-ER compartments, whereupon it is allowed for deglycosylation digestion and proteolytic processing. The resulting ^~^85-kDa glycoprotein of Nrf1D^ΔDC^ and its N-terminally-truncated ^~^75-kDa isoform have thus lost protection by membranes, such that both become more susceptible to protease attack, as compared to equivalents of the TMc-containing Nrf1.

### Nrf1D has a thiol-based redox-sensitive module in its unique TMc region

The above-described evidence demonstrates that the unique TMc region of Nrf1 dictates distinctions in the intrinsic regulation of this variant from wild-type Nrf1 and its mutant Nrf1D^ΔDC^. Together with bioinformatic analysis of distinct TMc regions in between Nrf1 and Nrf1 (Fig. 1, B to D), it is inferable that Nrf1D possesses a thiol-based (i.e. cystines 729, 730 and 738) redox-sensitive hydrophobic TMc module, that is topologically integrated within ER membranes (Fig. 5B). To examine this thiol-sensitive module, total lysates of COS-1 cells that had been transfected with anexpression construct for Nrf1, Nrf1, or Nrf1D^ΔDC^ were subjected to *in vitro* redox reactions with 5 mMol/L of hydrogen peroxide (H_2_O_2_) or dithiothreitol (DTT), respectively. These resulting reactants were analyzed by Western blotting with different antibodies against Nrf1 (Fig. 6A) or its C-terminal V5 tag (Fig. 6B). Of note, no obvious changes in the electrophoretic mobility of Nrf1 reactants with either H_2_O_2_ or DTT were examined herein. By sharp contrast, two longer Nrf1 isoforms of ^~^140- and 120-kDa (alongside with another ^~^75-kDa isoform) resolved on SDS-PAGE gels were obviously enhanced following oxidative reactions with H_2_O_2_, whilst reduction reactions with DTT led to significant increases in abundances of shorter molecular weight isoforms of between ^~^46-kDa and ^~^20-kDa (Fig. 6, A & B). However, much less of these smaller isoforms were detected in Nrf1-expressing cell lysates. Collectively, these demonstrate that various lengths of TMc-adjoining polypeptides are progressively truncated from within Nrf1, rather than Nrf1, in particularly reductive conditions. This notion is further supported by the converse evidence that almost none of similar shorter polypeptides were determined in Nrf1D^ΔDC^-expressing cell lysates, irrespective of whether they had reacted with H_2_O_2_ or DTT (Fig. 6, A & B). In addition, an extra polypeptide of Nrf1D^ΔDC^ with a heavier molecular mass of ~160-kDa was resolved by SDS-PAGE gels following its reactions with H_2_O_2_ but not DTT.

To determine whether the redox-sensitive TMc region of Nrf1D is thiol-site specific, three expression constructs for its mutants of triple, double or single cystines at positions 729, 730 and 738 into serine residues (i.e. Nrf1D^C729/730/738S^, Nrf1D^C729/730S^ and Nrf1D^C738S^) were made and then transfected into COS-1 cells, before redox reactions with H_2_O_2_ or DTT. Subsequently, Western blotting with Nrf1D-specific antibody revealed that almost all of other Nrf1D-derived isoforms between ^~^160-kDa and 25-kDa, except one ^~^75-kDa polypeptide, were substantially prevented by these two Cys-to-Ser mutants Nrf1D^C729/730S^ and ^Nrf1DC729/730/738S^ (Fig. 6C), but this seemed to be unaffected by redox reactions with H_2_O_2_ or DTT. Similar data were also obtained from anti-V5 antibody immunoblotting of Nrf1D^C729/730S^ and Nrf1D^C729/730/738S^ (Fig. 6D). This indicates that these polypeptides arising from both Nrf1D^C729/730S^ and Nrf1D^C729/730/738S^ are unstable so as be rapidly degraded, which results predominantly from Cys^729^/Cys^730^-to-Ser mutants. Further comparisons revealed that Nrf1D^C729/730/738S^, rather than Nrf1D^C729/730S^, conferred on two major Nrf1D-derived ^~^140- and ^~^120-kDa isoforms to be endowed with a slower electrophoretic shift to the locations of ^~^160- and ^~^130-kDa respectively (Fig. 6,C & D). Similar slower migration of small molecular weight isoforms between ^~^60-kDa and ^~^25-kDa was also examined in the case of Nrf1D^C729/730/738S^, but not of Nrf1D^C729/730S^ (Fig. 6D). From this, it is inferable that a possible proteolytic processing event is blocked by the Cys^738^-to-Ser mutant in the contexts of Nrf1D^C729/730/738S^. Overall, different results from Nrf1D^C729/730S^ and Nrf1D^C729/730/738S^, when compared with Nrf1D, demonstrate that the membrane-topological folding of Nrf1D, its post-translational modification and/or proteolytic processing are influenced by its Cys-to-Ser mutants within TMc (leading to dual decreases in its (x-helical hydropathicity and site-specific instability, as calculated in Fig. S6B).

**Figure 6.**
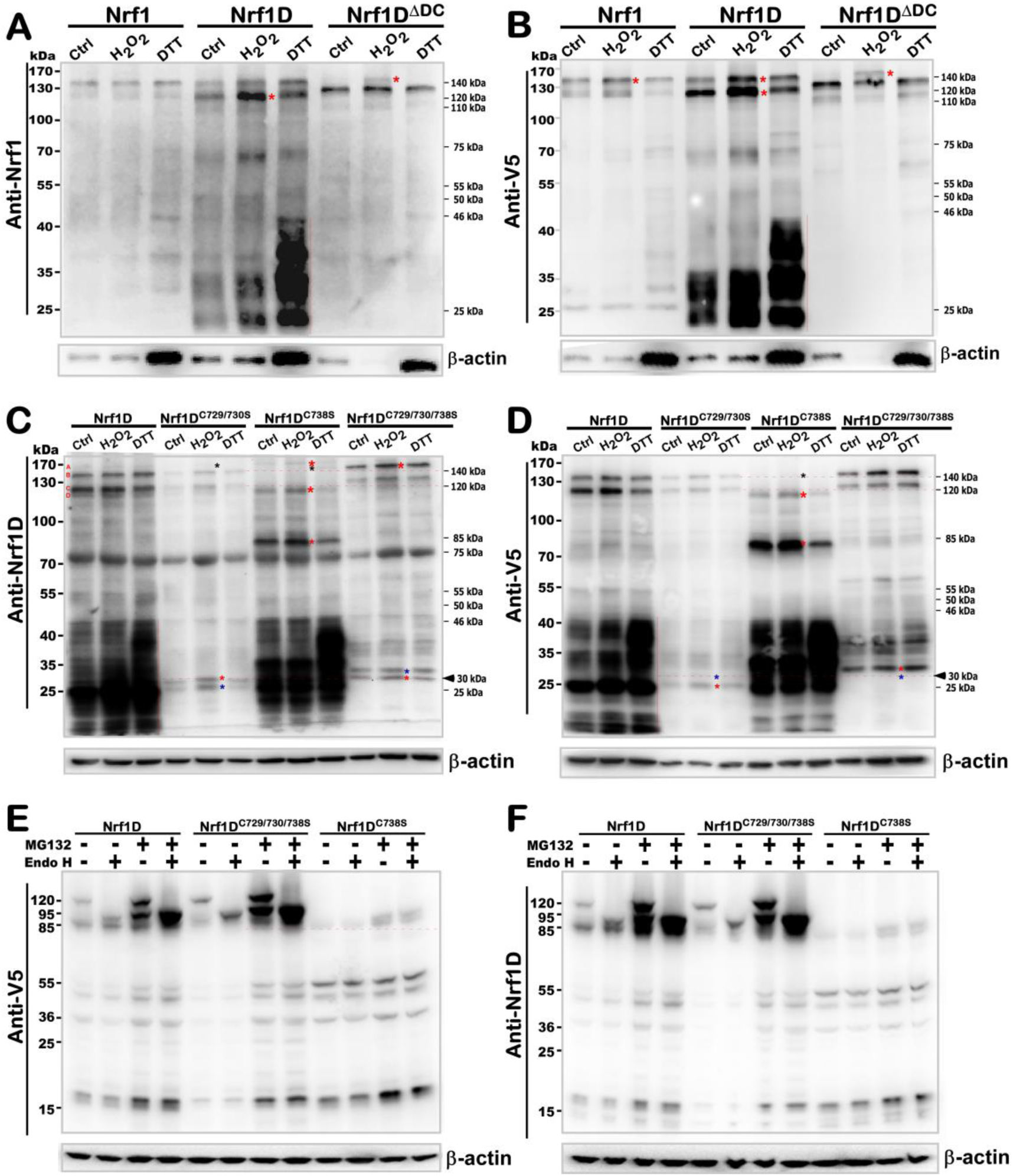
Redox reactions of Nrf1D and its mutants with H_2_O_2_ and DTT. (***A,B***) Total lysates of COS-1 cells that had been transfected with an expression construct for Nrf1, Nrf1D or Nrf1D^ΔDC^ were subjected to *in vitro* redox reactions with H_2_O_2_ (5 mmol/L) and DTT (5 mmol/L) for 10 min on ice, prior to be isolated by SDS-PAGE (with two-layers made of 8% and 12% polyacrylamide, respectively) and then examined by immunoblotting with antibodies against Nrf1 or V5 tag. (***C,D***) Similar redox Western blotting was employed for analysis of Nrf1D and its mutants (i.e. Nrf1D^C729/730S^, Nrf1D^C738S^ and Nrf1D^C729/730/738S^) that were incubated with H_2_O_2_ and DTT as described above. The redox reaction products were visualized by antibodies against Nrf1D or its C-terminal V5 tag. (***E,F***) COS-1 cells had been transfected with expression constructs for Nrf1, Nrf1D^C738S^ and ^Nrf1DC729/730/738S^ and treated with 5 μmol/L MG132 for 2 h before being harvested. The cell lysates were allowed for *in vitro* deglycosylation digestion by Endo H (+) or not (−). The resultant products were resolved by 4-12% LDS-NuPAGE gels and then determined by immunoblotting with indicated antibodies.

The above assumption is also supported by further experimental evidence, revealing that the single Cys^738^-to-Ser mutant (i.e. Nrf1D^C738S^) caused almost all of Nrf1D-derived ^~^140- and ^~^120-kDa isoforms to be completely abolished and then replaced by additional two minor polypeptides of ^~^160-kDa and ^~^115-kDa (Fig. 6C). The former ^~^160-kDa was not detected by immunoblotting with V5 antibody (Fig. 6D), implying that its C-terminally-tagged V5 ectope may be truncated. Surprisingly, changes of these longer isoforms were also accompanied by the emergence of an additional polypeptide of ^~^85-kDa arising from Nrf1D^C738S^, but not Nrf1D^C729/730S^ or Nrf1D^C729/730/738S^ (Fig6, C & D), suggesting that it is likely yielded from a potential proteolytic processing of unstable Nrf1D^C738S^-derived polypeptides. Interestingly, the existing longer isoforms of Nrf1D^C738S^ between ^~^160-kDa and ^~^85-kDa were further diminished by reactions with DTT, but not H_2_O_2_ (Fig. 6,C & D), indicating it become more unstable in the reductive conditions. By contrast, all those major polypeptides arising from Nrf1D^C729/730/738S^ and Nrf1D^C729/730S^ were modestly promoted by reactions with H_2_O_2_ and/or DTT (Fig. 6,C & D). Taken together with the aforementioned data (Figs. 5, S6 and S7), it is deduced that both Cys^729^ and Cys^730^ residues of Nrf1D are critical for TMc to serve as a thiol-based redox-sensitive module, whereas the adjacent Cys^738^ is essential for monitoring the membrane-topological folding of TMc, its dynamic repositioning, and proteolytic processing to distinct isoforms. The conclusion is further supported by the converse evidence obtained from Nrf1D^C738S^; this mutant led to no obvious alterations in the abundances of smaller isoforms between ^~^46-kDa and ^~^20-kDa yielded from the mutant protein processing and their electrophoretic mobility, when compared with those arising from Nrf1D, which were distinctive from the equivalents of Nrf1DC^729/730/738S^ (Fig. 6, C & D). However, it was found that increased abundances of those lower molecular-weight polypeptides of between ^~^46-kDa and ^~^20-kDa arising from Nrf1D and Nrf1D^C738S^, but not Nrf1D^C729/730S^ or Nrf1D^C729/730/738S^, were examined following *in vitro* reactions with DTT rather than H_2_O_2_. Collectively, such reductive reactions enabled Nrf1D^C738S^ to become more vulnerable (than Nrf1D) to its selective proteolytic processing, implying requirements of the Cys^738^ residue for the proper folding of Nrf1D and its stability.

Next, to clarify whether the processing of Nrf1D and its TMc mutant proteins is affected by 24-h incubation of COS-1 and HepG2 cells to a redox stressor *tert*-butylhydroquinone (tBHQ) or DTT, these cells were examined by Western blotting with three different antibodies against Nrf1β, Nrf1D or its C-terminally-tagged V5. The results demonstrated that both Nrf1D^C729/730/738S^ and Nrf1D^C729/730S^ gave rise to two major isoforms between ^~^140-kDa and ^~^120-kDa with slight slower mobility than equivalents of Nrf1D (Fig. S8, A to C). Such two isoforms were, however, completely abolished by a single point mutant Nrf1D^C738S^, but appeared to be replaced by additional two isoforms of ^~^115-kDa and ^~^105-kDa. The latter two major proteins of Nrf1D^C738S^ were electrophoretically migrated to similar locations of the N-terminally-truncated isoforms arising from Nrf1 or Nrf1D^C729/730/738S^, and also unaffected by deglycosylaton reaction with Endo H (Fig. 6E,F). Together with the above-described data (Fig. 6C,D), these suggest that the Nrf1D^C738S^ mutant proteins are rapidly processed to remove different lengths of its N-terminal portions so as to yield several proteoforms of Nrf1D^C738S^ between ^~^115-kDa and ^~^14-kDa. All these proteoforms arising Nrf1D^C738S^ and Nrf1D^C729/730/738S^, by comparison with those yielded from Nrf1D, were differently recognized by distinct antibodies against Nrf1β, Nrf1D or its C-terminally-tagged V5. These data indicate that the N-terminal and C-terminal portions of Nrf1D ^C738S^ and ^Nrf1DC729/730/738S^ are proteolytically processed in different ways. This notion is also further supported by additional evidence obtained from Western blotting of HepG2 cells that had been transfected with an expression construct for Nrf1, Nrf1DC^729/730/738S^ or Nrf1D^C738S^. In addition, mobility of most proteoforms was less or not influenced by treatment with tBHQ or DTT (Fig. S8, A to F), but this was strikingly distinctive from the aforementioned *in vitro* results (Fig. 6C,D). This distinction implies there exists a cytoprotective adaptation to redox stress.

### Nrf1D has only a low ability to transactive ARE-driven reporter gene induced by redox stress

To determine the Nrf1D activity to mediate ARE-driven reporter gene transcription, COS-1 cells were transfected with an expression construct for Nrf1D or Nrf1, together with *PSV40 nqo1-ARE-luc* and pRL-TK, and then treated with distinct redox inducers. As excepted, Figure 7A showed that Nrf1-mediated transactivation activity of ARE-Luc reporter gene was significantly induced to ^~^10-16-fold changes, as consistent with previous reports [22, 23], by Vitamin C (i.e. ascorbic acid, which has dual redox ability to donate electrons enabling it to act as a free radical scavenger and also to reduce higher oxidation states of iron to Fe^2+^; the Fe reducing activity enables hydroxyl radical to be produced, hence exerting a pro-oxidant effect [24]). However, only ^~^4-7-fold changes in Nrf1D-mediated reporter gene transactivation by ascorbate were determined (Fig. 7A), implying that Nrf1D has a lower activity than Nrf1 at mediating ARE-driven gene expression. Such lower activity of Nrf1D could be attributable to less abundance of two processed polypeptides similar to ^~^85- and ^~^75-kDa arising from the mature processing of Nrf1 (Fig. 7A, *lower panel*).

Further experiments revealed that Nrf1-mediated reporter gene was also substantially activated by stimulation of cells with either tBHQ or DTT, whilst Nrf1D only displayed a lower activity to transactivate the ARE-driven gene (Fig. 7,B & C). Intriguingly, treatment of cells with tBHQ or DTT caused a slightly faster electrophoretic migration of the full-length Nrf1D (of 749 aa) to ^~^110-kDa, only one longer polypeptide resolved by 4-12% LDS-NuPAGE gels, whereas intact Nrf1 (of 741-aa) was post-translationally processed to exhibit three or four major longer isoforms of between ^~^120-kDa and ^~^75-kDa (Fig. 7, B & C, *lower panels*). These findings indicate that the membrane-topological folding of Nrf1 and its proteolytic processing are certainly distinctive from those of Nrf1, particularly occurring under oxidative and/or reducing conditions. However, treatment of Nrf1-expressing cells with 12-*O*-tetradecanoylphorbol-13-acetate (TPA, a tumour promotor to induce AP-1 that comprises c-Jun and c-Fos) resulted in a significant decrease in the ARE-driven reporter activity (Fig. 7D). Similar decreases in the basal and Nrf1D-mediated activity of the reporter gene were also determined. In addition, no marked changes in the abundances of Nrf1- or Nrf1D-derived isoforms were examined following TPA treatment, albeit they appeared to be unstable ((Fig. 7D, *lower panels*). Together, Nrf1 (and/or Nrf1) -mediated ARE-battery gene expression is likely repressed by TPA-stimulated transcription factors (i.e. AP-1).

**Figure 7.**
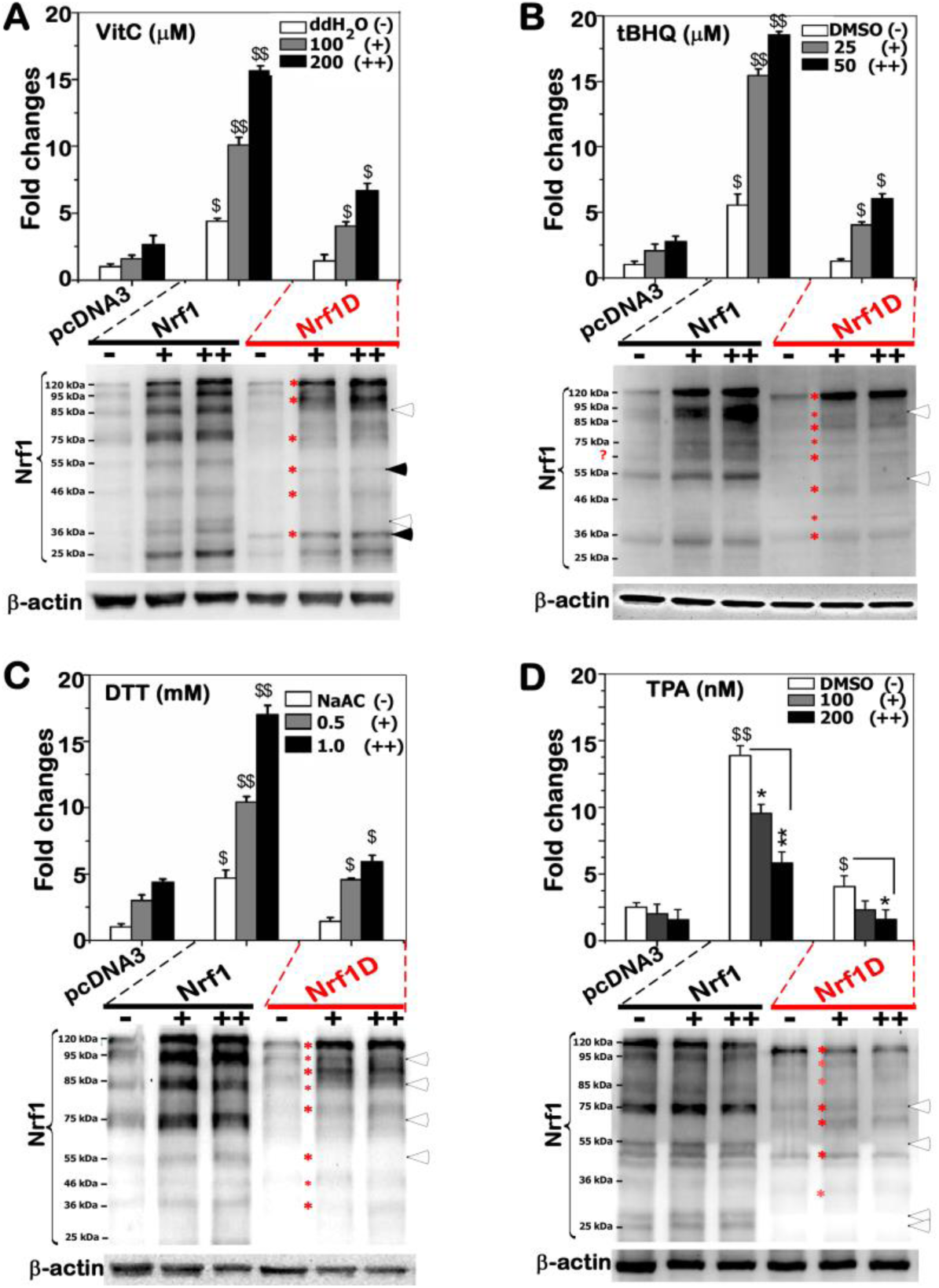
The activity of Nrf1D to regulate ARE-driven reporter gene in response to redox stress. COS-1 cells that had been transfected with an expression construct for Nrf1 or Nrf1, together with *P_SV40_Nqo1-ARE-*Luc and pRL-TK reporters (at a ratio of 10:5:1) were treated with indicated doses of VitC (***A***), tBHQ (***B***), DTT (***C***) or TPA (***D***) for 24 h before being measured. Subsequently, basal and chemical-stimulated ARE-driven reporter activity regulated by Nrf1 or Nrf1D was calculated as a ratio of its values against the background activity (i.e. measured following co-transfection of empty pcDNA3.1/V5 His B vector and ARE-driven reporter in response to indicated vehicle controls). Of note, the basal activity of empty vector was given the value of 1.0, and other data were calculated as fold changes (mean ± S.D) relative to this value. The data presented each represent at least three independent experiments, each of which were performed in triplicate, followed by statistical analysis of significant differences. Furthermore, the cell lysates were also resolved by 4-12% LDS-NuPAGE gels and examined by Western blotting with V5 antibody.

## DISCUSSION

In the present study, Nrf1 is identified as a candidate secretory transcription factor, with its precursor existing as a unique redox-sensitive transmembrane-bound CNC-bZIP protein detected in several somatic tissues (e.g. heart, lung, liver and testis, as well as bone borrow) before unleashed into the blood plasma. Collectively, the salient feature of Nrf1 is principally dictated by its C-terminal extra 80-aa region, allowing this variant protein to entail similar, but different, membrane-topology from that of prototypic Nrf1 (Fig. 8).

To give a better interpretation of Nrf1 topobiology, a model (Fig. 8A) is proposed on the basis of ever-accumulating evidence obtained from previous studies by us [9, 18, 25-27] and other groups [28-30]. Upon translation of Nrf1, its newly-synthesized polypeptide is targeted and anchored within ER membranes through its NHB1 signal peptide, spinning in an N_cyto_/C_lum_ orientation (i.e. its N-terminal and C-terminal ends face the cytoplasmic and luminal sides, respectively). The NHB1-adjoining portions of Nrf1 are successively translocated into the ER lumen, in which it is N-glycosylated by oligosaccharyltransferases to yield an inactive glycoprotein, but its DNA-binding (CNC and bZIP) domains are retained on the cytoplasmic side of membranes (Fig. 8A). When required for biological cues, some ER luminal-resident transactivation domain (TAD) elements will be dynamically repositioned through p97-fueled retrotranslocation into extra-ER cyto/nucleoplasmic sides of membrane, whereupon Nrf1 is allowed for deglycosylation by N-glycosidases and ubiquitination by Hrd1. Such modified Nrf1 protein is then subjected to the selective proteolytic processing to remove distinct lengths of its N-terminal portions by cytosolic DDI-1/2 protease-mediated cleavage and/or 26S proteasome-limited degradation. In turn, transcriptional expression of p97, DDI-1, and proteasomal genes is predominantly regulated by processed mature Nrf1 factor through coupled positive and negative feedback circuits [26, 27]. In addition, Nrf1 is also likely sorted out of the ER membrane and transported in the inner nuclear membrane to gain a direct access to target genes [25].

**Figure 8.**
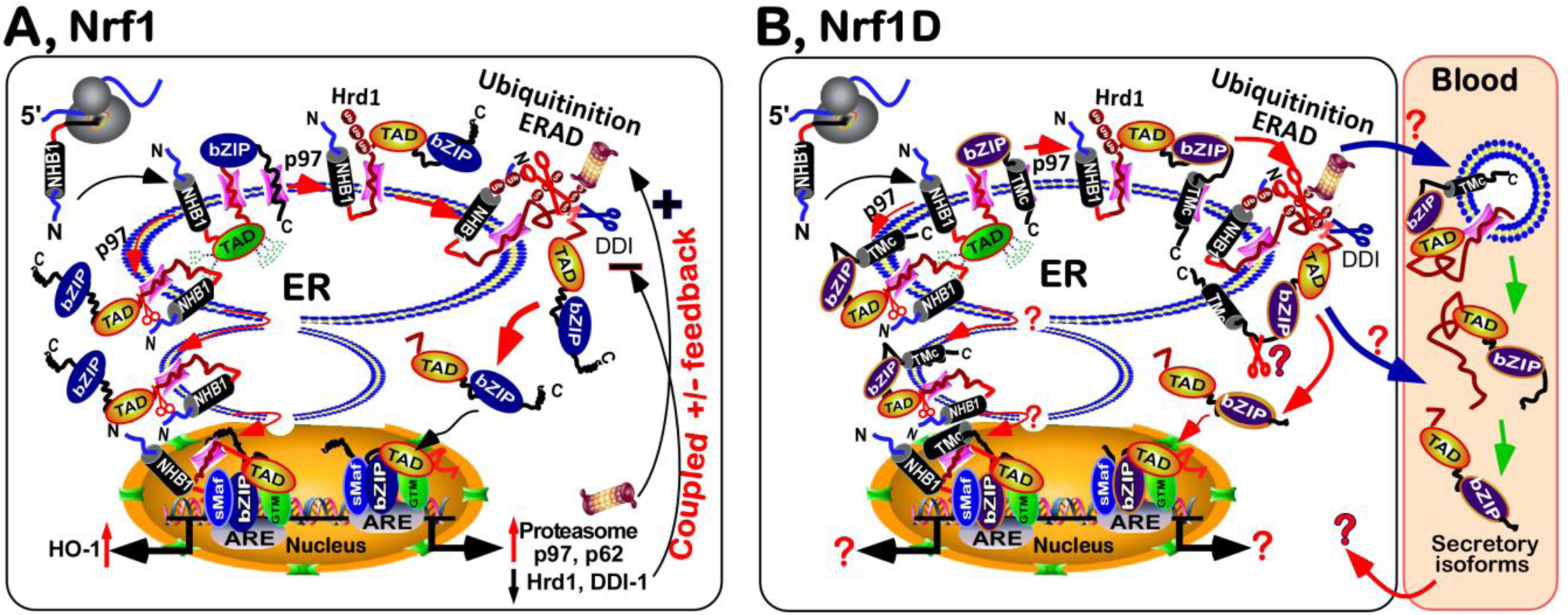
Two similar but different models proposed to explain distinctive topobiology of Nrf1D and Nrf1. (***A***) A model is proposed to interpret dynamic membrane-topologies of Nrf1 folded within the ER and dislocated to the nucleus. Of note, distinct topovectorial processes of Nrf1 dictates its post-translational modifications and selective proteolytic processing to yield multiple isoforms, as well as its transcriptional activity to mediate target gene expression. In turn, expression of Nrf1-target p97, DDI-1, and proteasomal genes is also predominantly regulated by processed mature Nrf1 factor through coupled positive and negative feedback circuits [26, 27]. (***B***) Another similar but different model is introduced to explain topobiology of Nrf1, that has a unique redox-sensitive TMc module spinning the ER membrane with an N_lum_/C_cyto_ orientation. Importantly, a few of Nrf1 isofroms are unlashed from the ER to enter the blood plasma (*right panel*), implying that it is the first candidate secretory transcription factor, albeit the detailed mechanisms remain to be further determined.

By contrast, Nrf1 is also a moving membrane-protein that entails dynamic topologies distinctive from prototypic Nrf1 (Fig. 8B). This distinction is determined by the unique C-terminal 80-aa region of Nrf1D, which substitutes the C-terminal 72-aa residues of wild-type Nrf1 (Fig. 1A). The C-terminal 80-aa region of Nrf1D is composed of a variant ZIP domain (that is certainly conserved with c-Fos and Fra-2, Fig. 1B) and another integral membrane-spinning TMc stretch (Fig. 1C). The variant bZIP of Nrf1D enables it to make distinctive dimerization from that made by the original bZIP of Nrf1, leading it to exert discrepant activity to mediate transcriptional expression of cognate target genes. However, the transcriptional activity of Nrf1 is tightly monitored by its topovectorial process, which is also dictated by its unique redox-sensitive TMc module (Fig. 8B), in addition to the NHB1 signal sequence, thus enabling it to be folded and processed in similar but different ways to account for Nrf1 (Fig. 8A).

In combination with our previous studies of Nrf1 [9, 18, 25-27], the evidence that has been presented here, reveals that Nrf1D is integrated in dynamic membrane-topologies within and around the ER (Fig. 8B), which are predominantly determined by both its N-terminal signal anchor (i.e. NHB1) and its C-terminal redox-sensitive TMc module. Within Nrf1, its amphipathic NHB1 peptide adapts an N_cyto_/C_lum_ orientation spinning membranes, whereas the hydrophobic TMc peptide is inferable to orientate in an N_lum_/C_cyto_ fashion across membranes. The acidic NST-containing TADs of Nrf1 are *cis*-translocated in the ER lumen allowing it to be N-glycosylated, whilst its basic aa-enriched CNC and bZIP domains are positioned on the extra-luminal cyto/nucleoplasmic sides of membranes. Once some TAD-adjoining portions of Nrf1 are repartitioned and retrotranslocated out of ER membranes, it should be allowed for deglycosylation and proteolytic processing by similar mechanisms accounting for Nrf1. However, the C-terminal redox-sensitive TMc module of Nrf1 enables it to exhibit its unique membrane-topological behaviour, such that this variant is also specifically processed in an additional way distinctive from the Nrf1 processing. The live-cell imaging of Nrf1D (C-terminally attached to GFP) demonstrates that the local membrane-topology of TMc, though having a higher hydrophobicity than the amphipathic NHB1 peptide, is still dynamically repositioned out of ER, so that it is disloacted into extra-ER cytoplasmic compartments, whereupon it loses the membrane protection against digestion by cytosolic proteases. Further experimental evidence unravels that the repositioning of Nrf1D and its selective proteolytic processing are under strict quality control by the precision thiol-based redox-sensitive module of TMc. For example, the Cys^738^-to-Ser mutation of TMc gives rise to an unstable protein of Nrf1D^C738S^, which could be allowed for fast dynamic repositioning into the extra-ER sides of membranes and rapidly proteolytic processing insomuch as that the putative glycoprotein and deglycoprotein disappear abruptly and is replaced by several processed isoforms distinctive from equivalents arising from Nrf1D (Figs. 6 & S8). By sharp contrast, the di-cysteine-to-Ser mutant Nrf1D^C729/730S^ leads to a slower electrophoretic mobility of glycoprotein and deglycoprotein when compared with those of Nrf1D. Such being the case, the redox-sensitive TMc of Nrf1D still enables it to be endowed with a less activity than wild-type Nrf1 to mediate ARE-driven reporter gene upon stimulation by redox stressors.

The low activity of Nrf1D is also attributed to a possible restriction of its juxtanuclear location detected in the bone marrow-derived cells (Fig. 3). Another reason is that it is, much to our surprise, discovered that some isoforms of Nrf1D are present in the blood plasma (Fig. 4). This exciting discovery demonstrates that Nrf1D has possibly secretory isoforms released through the proteolytic processing of this variant CNC-bZIP transcription factor to enter the blood plasma (Fig. 8B), albeit its precursor exists as a unique transmembrane protein in somatic tissues (Fig. 2). However, the detailed topovectorial mechanisms whereby Nrf1D is proteolytically processed to be secreted into the blood plasma remains elusive. For example, this is a very big difficult question to address where the intracellular Nrf1D is released and how it is delivered to the blood plasma. The plasma isoforms of Nrf1D may be derived from its founding mast cells and/or dendritic cells in the blood and other hematopoietic tissues [15], but non-hematopoietic tissues cannot also be ruled out.

### Concluding comments

Here, Nrf1D is identified as the first candidate secretory transcription factor, although it is largely unknown about the detailed mechanisms whereby its precursor transmembrane-bound CNC-bZIP protein is proteolytically processed in close proximity to the ER and unleashed into the blood plasma. Anyhow, this disruptive discovery opens a whole new avenue for gene regulation by Nrf1D secreted from cell communication (even from blood transfusion). However, this work also raises a lot of new questions to attract wide attentions from researchers in all relevant fields. In addition, we should notice that inhibitor of DNA binding 1 (ID1) was determined as a basic helix-loop-helix (bHLH)-ZIP transcription factor and secreted only in the synovial fluid, but not the blood plasma, of model animals with rheumatoid arthritis [31, 32]. However, this small bHLH-ZIP protein does not possess any of the putative transmembrane domains, similar to those within Nrf1D.

## Materials and Methods

### Chemicals, antibodies and plasmids

All chemicals were of the highest quality commercially available. Amongst them, proteinase K (PK), Digitonin and Endo H were purchased from New England Biolabs, whereas MG132, CHX, DMSO, VitC, tBHQ, TPA and DTT were obtained from Sigma, USA. Mouse monoclonal antibodies against the V5 epitope was from Invitrogen Ltd. Of note, two specific antisera against Nrf1 or Nrf1D were produced in rabbits by our laboratory in collaboration with commercial companies. Two expression constructs for Nrf1 and ER/DsRed had been described previously [14, 25]. Here, Nrf1D was engineered by inserting the XbaI/SacII fragment (encoding aa 1–669 of Nrf1) with an Nrf1D-specific cDNA sequence (encoding its unique aa 670-749, which was synthesized by a commercial company) into the pcDNA3.1/V5-His B vector. Additional 3 cysteine-to-serine mutants Nrf1D^C729/730/738S^, Nrf1D^C729/730S^ and Nrf1D^C738S^ were created separately. The C-terminally-truncated Nrf1D^ΔDC^ was also made by only inserting the cDNA sequence encoding the N-terminal 669 aa of Nrf1. Other expression constructs for two fusion proteins Nrf1D-GFP or Nrf1D^ΔDC^-GFP were generated by ligating their encoding sequences to the 3-end of GFP-encoding cDNA into the KpnI/BstBI site of pcDNA3.1/V5-His B, respectively. The fidelity of all cDNA products was confirmed by sequencing.

### Organ collection and blood fractionation

All of experimental organs and blood samples were collected from 8-weeks-old male or female BALB/c mice. After these mice were narcotized by diethyl ether, the blood samples were obtained from their eyeballs, that were perfused with PBS containing 10 mU/ml heparin. The bloods were fractionated by centrifuging at 2000 rpm for 10 min for a collection of upper layer (i.e. blood plasma), middle layer containing white blood cells (WBCs), and lower layer containing red blood cells (RBCs). After WBCs and RBCs were rinsed by a serum-free medium, they were collected by centrifuging at 2500 rpm for 5min (which were repeated for 3 times), prior to being stored at -80°C for the further use. Thereafter, the mice were sacrificed by being decapitated to obtain relevant tissues and organs, which were harvested and immediately snap-frozen in liquid nitrogen for further analyses. In addition, the bone marrow was also obtained from the tibiae and femurs of these mice by washing with PBS. All these animal experiments were indeed conducted according to the valid ethical regulations that have been approved.

### Real-time qPCR analysis

Total RNAs were extracted from examined tissues and then reverse-transcripted to yield the first single-stranded cDNAs, which served as temples of real-time quantitative PCR, according to the manufacturer’s protocol (TaKaRa, Dalian, China). To investigate differential expression of both Nrf1D (Genbank No. AF071084.1) and Nrf1 (Genbank No. NM_008686.3) at their endogenous mRNA levels, a pair of optimal primers complementary to the indicated upstream (5’-GCTGACTTCCTGGACAAGCAGATGA-3’) and downstream (5’-GGAACCAACCACCTGGCTATC-3’) nucleotide sequences was designed and then added in the PCR. The reaction was conducted in the following conditions: inactivation at 94°C for 2 min followed by 34 cycles of 10 sec at 98°C denatured, 30 sec at 58°C annealed, and 2 min at 58°C extended, with an additional extension of 7 min before being stopped. The resulting PCR products were isolated by 1% agarose gel electrophoresis and collected for cDNA sequencing. In addition, expression of β-actin or RPL13A mRNAs was used as an internal control for normalization.

### Fluorescence *in situ* hybridization (FISH)

The bone marrow cells were fixed in 10 ml of 4% paraformaldehyde for 15 min at 4°C, and pelleted by centrifuging at 1000 rpm for 5 min. After the supernatants were abandoned, the cells were permeabilized for 10 min by 0.1% Triton X-100 at normal temperature, and then collected by centrifuging at 1000 rpm for 5min. The cells were adjusted to an appropriate concentration of 3×10^6^/ml of PBS and dropped on glass slides. Subsequently, the slides were immersed in a 2× SSC solution at 37°C for 5 min, placed in a proper order of 70%, 85% and 100% ethanol at room temperature for 3 min each for dehydration, and then dried naturally. A hybridization solution was preheated at 58°C for 2 h, and then added to the above slides at a total volume of 10 μl, which contained 1 μl of each probe of Nrf1 (5’-CGCACGATGGCTGA CCAGCAGGCTC-3’, which retained within Nrf1 but lacked in Nrf1D, with its 5’-end labeled by 5-FAM, emitting a green fluorescence) and Nrf1D (5’-CGATTGCTTCGAGAAAAGGAAAATGAGAAGTGC-3’, which was labeled by Texas Red at its 5’-end so as to emit a red fluorescence). These probes were allowed to hybridize with indicated samples overnight at 37°C in a hybridization chamber. The probe-hybridized sample slides were washed in 0.4× SSC plus 0.3% Tween-20 at 65°C for 3 min, followed by 2× SSC plus 0.1% Tween-20 at room temperature for 30 min, and then stained with Wright-Giemsa and DAPI, respectively. All probes used in this study were synthesized by Invitrogen (Shanghai, China).

### Cell culture, transfection, and reporter gene assays

Unless otherwise indicated, monkey kidney COS-1 cells (3×10^5^, which had been previously purchased from ATCC and maintained in our laboratory) were seeded in 6-well plates and grown for 24 h in Dulbecco’s Modified Eagle Medium (DMEM) containing 10% fetal bovine serum (FBS). After reaching the 70% confluence, the cells were transfected with a Lipofectamine 3000 (Invitrogen) mixture that contained an expression construct for Nrf1, Nrf1D or mutants, together with two reporters *P_SV40_Nqo1-ARE*-Luc and pRL-TK (at a ratio of 10:5:1). The latter reporter was used as an internal control for transfection efficiency, whilst additional mutant version of these reporter genes that lacked ARE sequences served as a negative control. At 24 h after transfection, the cells were further treated by different doses of DTT, tBHQ, VitC or TPA for an additional 24 h. Subsequently, ARE-driven luciferase reporter activity and western blotting were measured approximately 24 h after transfection alone or plus chemical treatments. Basal and stimulated reporter gene activity regulated by Nrf1, Nrf1D or mutants was calculated as a ratio of its values against the background activity (i.e. measured following co-transfection of empty pcDNA3.1/V5 His B vector and ARE-driven reporter in cellular response to indicated vehicle controls). Of note, the basal activity of empty vector was given the value of 1.0, and other data were calculated as fold changes (mean ± S.D) relative to this value. The data presented each represent at least three independent experiments, each of which were performed in triplicate, followed by statistical analysis of significant differences.

### Live-cell imaging combined with *in vivo* membrane protease protection assays

The live-cell imaging was performed as described by [8, 9]. COS-1 cells (2×10^5^) were seeded in 35-mm dishes and cultured overnight. The cells were then co-transfected for 6 h with 3 lag DNA of an expression construct for Nrf1D-GFP or Nrf1D^ΔDC^-GFP and 0.5 lag DNA encoding ER/DsRed (as an ER luminal-resident protein marker [14, 18, 33]). Subsequently, the cells were allowed for a 16-h recovery from transfection, the plasma membranes of COS-1 cells were permeabilized by digitonin (20 μg/ml) for 10 min. Thereafter, the cells were subjected to *in vivo* membrane protection reactions against digestion by PK (50 μg/ml) for 35 min. During the experiment, live-cell images were acquired every minute on Leica DMI 6000 green and red fluorescence microscopes equipped with a high-sensitivity HAMAMATSH ORCAER camera. Relative fluorescence units were measured by using the Simulator SP5 Multi-Detection system for GFP with 488-nm excitation and 507-nm emission, and for DsRed2 with 570-nm excitation and 650-nm emission.

### Deglycosylation assay, western blotting and Coomassie brilliant blue staining

Experimental cell lysates are subjected to *in vitro* deglycosylation digestion by Endo H, according to the manufacturer’s instruction (New England Biolabs, USA). Equal amounts of proteins prepared from cell lysates were loaded into each of electrophoretic wells, and visualized by Western blotting with distinct antibodies against Nrf1, Nrf1D or its C-terminally-tagged V5 ectope. β-Actin served as an internal control to verify amounts of proteins loaded in each well. On some occasions, nitrocellulose membranes that had already been blotted with antibodies were rinsed for 30 min with a stripping buffer before being re-probed with additional primary antibody [34]. Furthermore, some protein-blotted gels or nitrocellulose membranes were stained by 2.8 μg/ml of Coomassie brilliant Blue R-250 in 10% acetic acid. After the protein bands appeared in 30 min, the gels were de-stained in 10% acetic acid for over 2 h.

### Immunoprecipitation and immunohistochemistry

Blood samples were extracted in RIPA lysis buffer (Beyotime, China) and then subjected to immunoprecipitation with Protein-G Sepharose 4 Fast Flow beads, after incubation with specific antibodies against Nrf1 or Nrf1D at 4°C overnight. The Protein-G Sepharose beads were washed 3 times with PBS. The resulting immunocomplexes with Nrf1 or Nrf1D antibodies were visualized by immunoblotting with another indicated antibodies. For immunohistochemistry analysis, distinct tissues were incubated with the primary antibodies against Nrf1 or Nrf1D and the secondary anti-IgG (BIOPIKE, USA), according to the instructions of manufacturer.

### Statistical analysis

The significant differences in changes of Nrf1- or Nrf1D-mediated reporter gene activity, and the abundances of their transcript and protein expressed in different tissues were determined using the *Student’*s T-test or Multiple Analysis of Variations (MANOVA). The data are herein shown as a fold change (mean± S.D), each of which represents at least 3 independent experiments undertaken on separate occasions that were each performed in triplicate (n=9).

## Author contributions

J.Y., H.W. and S.L. performed most of experiments, collected the resulting data and prepared draft of most figures and supplementary information presented in this manuscript. Y.C. done mutagenesis of Nrf1D and time-course analysis. S.H. carried out immunoprecipitation. M.W. and L.Q. helped together with molecular cloning to create an expression construct for Nrf1D, animal experiments with organ collections, transfection and luciferase assays. Y.Z. designed this study, analyzed all the data with some bioinformatic tools, helped to prepare all figures, wrote and revised the paper.

## Acknowledgments

The study was supported by the National Natural Science Foundation of China (key programs 91129703, 91429305 and project 31270879) awarded to Prof. Yiguo Zhang (University of Chongqing, China), and in part funded by Chongqing University postgraduates innovation project (No. CYB15024) awarded to Mr. Lu Qiu.

